# A model for sheath formation coupled to motility in *Leptothrix ochracea*

**DOI:** 10.1101/157073

**Authors:** James Vesenka, James Havu, David Emerson

## Abstract

Optical and atomic force microscopy (AFM) of naturally occurring *Leptothrix ochracea* was used to study the fine structure of sheaths and cells. Morphology of young sheaths suggests the scaffold chains have strong self-adhesion. Evidence from un-encapsulated cells indicates fresh scaffold production through cell walls. Simple diffusion arguments are used to explain the morphology of the sheath structure. We propose a novel cell motility model based on previously published video data, our AFM images of naked cells, and simple flow calculations. The model indicates that motility results from differential shear forces resulting from extrusion of sheath material that passively pushes a filament of connected cells forward as the surrounding sheath material hardens behind the cell train.

## Introduction

*Leptothrix ochracea* is a common, sheath-forming microorganism that lives in freshwater habitats that have high concentrations of soluble, reduced iron (Fe(II)). Since iron is the 4^th^ most abundant element in the Earth’s crust, these habitats are common. They are typically found where there is standing or slowly flowing water enriched in Fe(II), examples include streams, wetlands, and springs, as well as technical environments like water distribution pipes (Emerson et al., 2010). Where these bacteria occur, it is common to find rust-colored deposits or precipitates of Fe-oxyhydroxides that form loosely aggregated microbial mats made up of a consortia of microorganisms, that are dominated by bacteria involved in Fe-cycling (Roden et al., 2012). When present, *L. ochracea* is often the dominant morphotype observed in these habitats due to its copious production of tubular sheaths that are encrusted with Fe minerals, principally ferrihydrite. These microbial iron mats can accrete rapidly, and due to their large surface area, the sheath/mineral matrix has the ability to influence the chemistry of the local waters that they contact (Chan et al., 2016).

While *L. ochracea* was first described 120 years ago, it has yet to be cultivated reliably in the laboratory, thus much about its life remains a mystery (van Veen, et al 1978). Recently, using cultivation-independent techniques it was shown that *L. ochracea* is a close relative of other sheath-forming species like *Leptothrix cholodnii* and *Sphaerotilus natans* (Fleming et al., 2011). Both of these latter organisms are heterotrophic bacteria that utilize organic matter for growth, and do not require Fe(II) as an energy source. *L. ochracea* does require Fe(II) for growth, although it remains unknown if Fe(II) is its primary energy source; nonetheless, it will not grow on the heterotrophic medium utilized by *L. cholodnii* (Fleming and Emerson, manuscript in preparation). Another important difference between these organisms, is that in cultures of *L. cholodnii,* the sheaths are nearly all full of cells, while for *L. ochracea,* even in the most actively growing microbial mats, only about 10% of the sheaths contain cells (van Veen, et al 1978; Chan, et al 2016). In all these organisms, sheath production is a coordinated event that results in the formation of a resilient biological structure. A sheath affords these microbes certain benefits, like protection from predators, primarily protozoa that cannot tear open the sheaths, and positioning, if the sheath is attached by a hold fast to a solid substrate, then the organism can maintain position in an environment favorable for its growth. In the case of *L. ochracea* it is also likely that the sheath plays an important role in protecting the cells inside from becoming encrusted in solid Fe-oxides that precipitate as a result of their growth.

*L. ochracea* is capable of prodigious sheath production: it is estimated that a chain of 100 cells that are about 0.5 mm in length can produce 1 meter per day of sheath (Fleming and Emerson, manuscript in preparation). The sheath is composed largely of Fe-oxyhdroxides that are by byproducts of Fe-oxidation. Analysis of *L. ochracea* sheaths collected from an iron mat by scanning transmission X-ray microscopy (STXM) and near edge X-ray absorption fine structure spectroscopy (NEXAFS) revealed that it contains an organic component, possibly a polysaccharide (Chan et al 2009). Consistent with these observations, Suzuki et al. have (2011) used high-angle annular dark-field scanning electron microscopy to show that *L. ochracea* sheaths contain iron, carbon and oxygen, again indicative of the sheath being composed of an organo-metallic matrix. However, beyond these physical observations little is known about the chemistry of the sheath, or its exact role in Fe-oxidation.

Much more is known about sheath formation in the heterotroph, *L. cholodnii.* The sheath of this organism consists of a fibrillar matrix, with individual fibers composed of complex polysaccharides made up of uronic acids, amino and neutral sugars, and peptides (Emerson and Ghiorse, 1993, Takeda et al., 2005). A major building block of these fibrils is a novel glycoconjugate that contains N-acetyl-L-cysteinylglycine. Takeda et al. (2009) have proposed that the sheath fibrils of *L. cholodnii* can spontaneously assemble through the formation of disulfide bonds from side chains containing L-cystine and hydrophobic interactions from 3- hydroxypropionic acid substituted for glycan units in the chain. The extensive disulfide bonding knits the fibrils together into a robust tubular structure that is resilient to chemical or physical disruption. Similar properties are shared with another sheath-former, *Sphaerotilus natans,* that is in the same family of microbes as *L. cholodnii* and *L. ochracea* (Takeda et al., 2007).

In addition to having fascinating chemical and physical properties that contribute to the unusual lifestyle of these bacteria, it is proposed that the nano-crystalline arrays of reactive iron-oxides that coat the *L. ochracea* sheath may have properties suitable for commercial applications (Kunoh et al., 2015). Some potential uses include serving a catalyst for organic synthesis (Ema et al., 2013), or improving the performance, cost, and life cycle of lithium-ion batteries (Sakuma et al., 2014). In addition, their capacity for adsorption of other metals like arsenic and lead, as well as organics could be useful in water treatment schemes that promote their growth using naturally Fe-rich waters (Emerson and de Vet, 2015).

Despite the many fascinating questions about sheath production in *L. ochracea,* the lack of a sufficient laboratory model for *L. ochracea,* means there is still much to learn about the mechanism of sheath formation or details of its ultrastructure. The work described here utilized atomic force microscopy (AFM) to gain a better understanding of the structural details of naturally occurring *L. ochracea* sheaths. The goal is to develop a model that explains their physical characteristics and possible mechanism of motility consistent with what is known about the biology of this unique microorganism.

## Methods

### Sampling for *L. ochracea* and epifluorescence imaging

Fresh sheaths of *L. ochracea* were collected form iron-rich seeps in Boothbay Harbor and Kennebunkport, Maine. The Boothbay site has been described previously (Chan et al 2016). The site in Kennebunkport (43.362° N, 70.477° W) was a small roadside ditch (pH 6.2) located beside a pine forest. This site had slowly flowing water 5 -10 cm deep. We were unable to measure the concentration of Fe(II) at this site. Nonetheless, the abundant presence of iron mats observed as flocculent, rust colored material that loosely adhered to the sediment is consistent with micromolar concentrations of Fe(II). Concentrated sheath samples in their original aqueous media was collected with a pipet into Eppendorf tubes and refrigerated at 4°C until being further processed, within 24 hours, as described below.

Cells within sheaths were located by staining and imaging by epifluorescence microscopy. Dilute solutions of Syto13 DNA stain (Molecular Probes/Invitrogen) were added (final Syto13 dilution 1:250) to these samples and samples were gently vibrated to disperse clumps of sheaths. 20μL aliquots of the solution were pipetted onto freshly cleaved mica and allowed to dry under ambient conditions – effectively incubating the stain with the cells for about 20 minutes while slowly drying. A Zeiss AX10 fluorescence microscope illuminated by Lumen Dynamics X-Cite Series 120Q UV source was used to identify regions containing intact sheaths, collapsed sheaths and sheaths containing *L. ochracea* cells (Fig 1). Fluorescent and bright field images were digitally collected to aid in "optical navigation" in the AFM. That is, once stained regions of interest were identified, the samples were transferred to the AFM and identified using the AFM's video microscope in order to place the AFM probe over the region of interest. The samples were imaged with a Digital Instruments Nanoscope IIIa controller and Multimode TopView AFM with long working distance (35mm) 400x optical navigation system in ambient air. No noticeable contamination (loss of resolution) of the tip when imaging sheaths was observed during imaging – the same tip could be used for days. However imaging naked cells did lead to tip contamination. The AFM probes used were Budgetsensors Single HiRes150 (resonance frequency ≈ 150kHz) to image the samples at a scan rate of 500nm/second and later analyzed using Nanoscope software.

**Figure 1:**
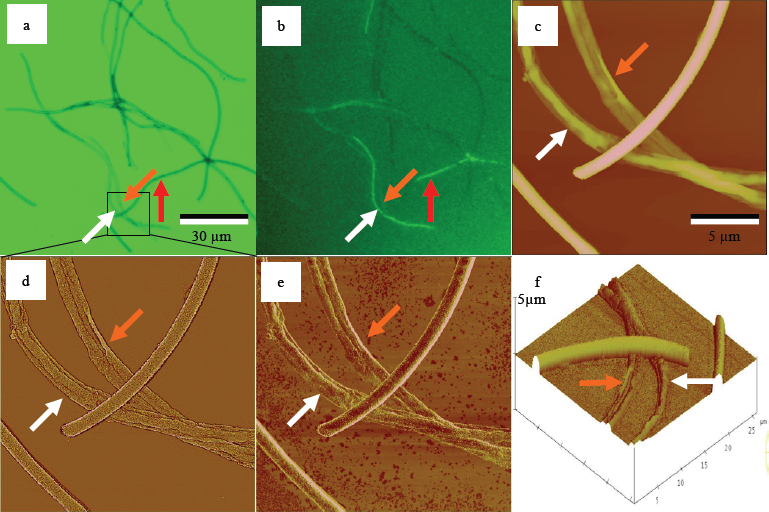
(a) Bright field image of intact and collapsed cells (b) and same region fluorescently stained Syto 13 revealing *L. ochracea* cells in sheaths. Red arrow labeled darker intact *L. ochracea* sheath with cells inside but invisible to the AFM because of the lack of topographic relief. Orange arrows represent empty collapsed sheath and white arrows identifies cells the AFM can detect below a collapsed sheath (c) AFM height image zoomed into boxed area in the bright field image and confirmed the state of the sheaths as either being intact or collapsed as well as the presence of the cells inside the collapsed sheath. (d) is a high-pass filtered height image used throughout the remainder of the figures to identify detailed nanostructures. (e) is a simultaneously captured phase image indicating adhesive interactions between tip and substrate but no obvious interactions with the sheaths. No obvious phase interactions were detected so only high passed filtering images are presented in the remainder of this paper (f) is a three-dimensional rendering of the height image clearing showing the substantial height differences between intact and collapsed sheaths. Optical images (a) and (b) are 150μm a side. AFM images (c)-(f) are 25.6μm scan size. Vertical scale on (c) is 0nm to 2000 nm, (d) is 0nm to 500nm, (e) is 0° to 50° (from dark brown to white). (f) has a highly exaggerated vertical scale of 5μm from the substrate to top of scale bar to more easily display differences between intact and collapsed sheaths clearly indicating the topographic nature of (c). The white and orange arrows identify the same features found in the other five in this rotated image.

## Results

### AFM Imaging

Fresh sheaths of *Leptothrix ochracea* from natural seeps were characterized using both optical and atomic force microscopies (Figure 1). A combination of brightfield and fluorescence imaging of Syto-13 stained samples was used to rapidly identify cells, as well as intact and collapsed sheaths for AFM imaging. It was routinely observed that the drying process required for AFM analysis resulted in immature sheaths, also referred to as "proto-sheaths", to collapse, and in cases where cells were present in the collapsed sheath they could be observed by AFM. This was fortuitous, since AFM is a surface sensing technique and unable to detect cells in intact (un-collapsed) sheaths. Brightfield (Figure 1a) and fluorescence (Figure 1b) microscopy made it possible to quickly identify intact (dark) and collapsed (fainter) sheaths and cell filaments stained by Syto13. The low magnification optical images were used to navigate to regions of interest for higher resolution AFM imaging. Figure 1c, d, and f is the region identified in the small square in in Figure 1a. Figure 1c is a top view image in which color indicates height above substrate, shown slightly rotated to enhance contrast in 3D in Figure 1f. Figures 1d, e are filtered images of Figure 1c to highlight the sheath ultrastructure. White arrows identify cells inside a collapsed sheath, orange arrows indicate an empty sheath, and the red arrows indicate cells inside an inaccessible (to AFM) intact sheath. Cells imaged inside sheaths appear twice as thick as the naked cells, which may be a result of protection from dehydration.

The use of high pass filtering (Figure 1d) and to a lesser extent phase imaging (Figure 1e) revealed details at nanometer resolution about the sheath and cell structure. Consequently, high pass filtering was used extensively for this work to characterize the irregular fibril-rich matrix of the sheath exterior (Figure 1d). Phase imaging can indicate tip-sample chemistry, but Figure 1e shows little contrast, which indicates the AFM probe did not adhere significantly to the sheaths. This is important because it ensured that sample details were not being obscured by tip-sample interactions and minimized imaging artifacts. Figure 1f shows a three-dimensional rendering of the intact and collapsed sheaths with a highly exaggerated vertical scale to emphasize the differences between collapsed and intact in sheath features.

Figure 2 provides examples of individual, intact fresh sheaths with a variety of internal fiber widths. The width of the individual fibrils on the mica surface averaged about 22±5nm (e.g. as seen in Figure 2b red arrow). These fibrils were about the same dimensions as the width of single fibers found within immature sheaths (e.g. Figure 2c, 24±5nm white arrow on sheath). This collapsed immature sheath also had a single wall thickness of 22±3nm (image not shown), suggesting it consists only of a single fiber layer. Figure 2c,f are examples of younger cell-filaments within new intact sheaths. Some intact sheath fibers (Figures 2d,e) and many fibers on collapsed sheaths had slightly larger diameters (35±10nm). Lastly, intact sheaths with much wider fibers averaging about 62±5nm in diameter were also observed, presumably these are more mineralized due to continued aquistion of Fe-oxides (e,g. Figure 2f). In Figures 2d-f note that the scaffold appears slightly oriented in the direction diagonal or parallel to the long length of the sheath. Details of numerous cell measurements taken by the AFM are summarized in Table 1.

**Figure 2:.**
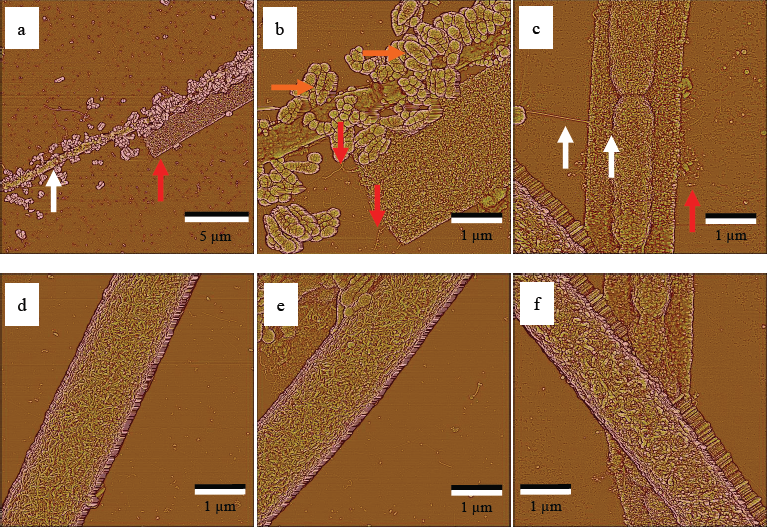
High pass filtered images immature sheaths and fresh intact sheaths have sharply defined edges. (a) is a low magnification view of naked cells (white arrow) and empty immature sheath (red arrow). (b) zooms into the center of (a) and highlights long linear sticky fibrils (red arrows, single fibrils of average width 22±5nm), also in c) that compose the scaffold of the sheaths. The purple arrows identify large, unidentified bulbous structures surrounding the cells, which appear to be associated with degradation of the cells. These structures are not found near the cells still encased in the sheaths as in the example shown in (c). Evidence that the cells are inside the sheath comes from the continuity of the scaffold over the mica substrate and cells (white arrow on sheath of average width 24±5nm). (d-f) exemplify how the fibrils are woven into the scaffold. In top view (not shown) the collapsed sheath is 45±3nm thick (top and bottom layer) and 22±3nm thick just the bottom layer. This suggests the sheath's consists of monolayers of fibrils. The color scale represents height out of the plane of the paper that is 0nm (dark brown) to 50nm (white) for all images.

**Table 1.** summarizes several hundred measurements as a set of ranges that appear to depend only on the age of sheaths, based on the observation that older sheaths have thicker walls. The outer diameters of the sheaths are a maximum of 1200nm with wall thicknesses ranging from 22 to 200 ±3nm depending on age of sheath. Details of the polysaccharide scaffold can be imaged at high resolution and suggest the sheath material is subject to strong self-adhesion. Lastly, high-resolution images of cells indicate the dried cells have diameters of approximately 500±10nm and lengths ranging from 1800 to 4100nm.

| Height<sub>cell</sub><br>(nm) | Width<sub>cell</sub><br>(nm) | Length<sub>cell</sub><br>(nm) | Thickness<sub>sheath</sub><br>collapsed (nm) | Width<sub>sheath</sub><br>collapsed (nm) | Height<sub>sheath</sub><br>intact (nm) | Width<sub>sheath</sub><br>intact (nm) | ID/OD<sub>sheath</sub><br>intact (nm) |
| --- | --- | --- | --- | --- | --- | --- | --- |
| 140-340 | 750-910 | 1800-4100 | 22 - 250 | 1800 - 2200 | 950 - 1200 | 1350 - 2200 | 970/1150 |

Figure 3 appears to be a fortuitous observation of a line of cells both partially within a fresh sheath and outside the sheath. Close examination of the naked cells indicates they are in the process of exuding sheath material (orange arrows surrounding cells in Figure 1c), perhaps at the instant the cells were drying down on the substrate. If this is the case we can surmise that the sheath material is exuded along the entire length of the cell out to a width approximately equivalent to that of the sheath dimensions. The material coming from these naked cells appears to be sticky and similar in properties to those in Figures 2b,c. This was confirmed by the observed loss of resolution while imaging naked cells that implies a different chemistry between tip and sample. Careful observation of the leading edge of the cells (Figure 3a white arrow), identified as lacking an obvious sheath (Figure 3a red arrow) of the cell line indicates the extruded material is thinner in the front of the chain than at the trailing edge, suggesting an accumulation of sheath material along the length of the cell filament.

**Figure 3:**
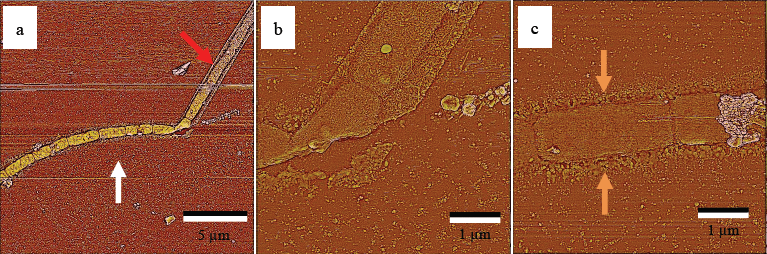
Overview (a) of high pass filter from topography of cells found both within (red arrow) and outside (white arrow) a fresh sheath. Within the sheath (b) the source of the sheath material coming from the cells is not possible to determine. However, when the cells are naked (c) scaffold material clearly appears to be exuded from the cell perimeter (orange arrows covering the length of both sides of the cell). The highest concentration of scaffold material appears to drop off with distance perpendicular to cell train direction to a width approximately equivalent to the sheath diameter seen in (b), suggesting a nascent stage of sheath production. The color scale represents height out of the plane of the paper that is 0nm (dark brown) to 50nm (white) for all images.

Figure 4a,b highlights the large differences in fibril lengths, confirmed by grain size analysis (Figure 4c,d). Figure 4a shows an immature collapsed around a cell filament on the left, and an older, intact sheath on the right; the vertical scale is exaggerated to show the height differences between collapsed and uncollapsed sheaths. The boxed region of Figure 4a is analyzed for grain size in Figures 4c,d. Grain size analysis consists of manually determining a background level that converts the area to be analyzed into a binary mixture of either bright islands, or "grains" consisting of contiguous pixels, separated by dark boundaries. The manual background (dark boundaries) determination consisted of visual identification of fibers that appeared to stick out from the surface of the sheath. Once the background level is determined the software performs a distribution analysis of grain frequency versus contiguous pixel area. Both small (Figure 4c) and large (Figure 4d) grain distributions were manually identified. Figure 4c records grains smaller than 4300nm^2^ (average 620nm^2^ with standard deviation of 770nm^2^); note how the smaller red highlighted polymers predominate over the immature sheath on the left side of the image. Figure 4d records grains larger than 4300nm^2^ (average of 7900nm^2^ with standard deviation 5400nm^2^); note how the longer fibrils predominate over the intact sheath to the right of the image. Note as well that the longer fibers are oriented more parallel to the sheath axis, as described above for Figure 2 d,e,f, and also shown in Figure 4b. We speculate that these differences are due to variations in growth conditions discussed in Figure 5.

**Figure 4:**
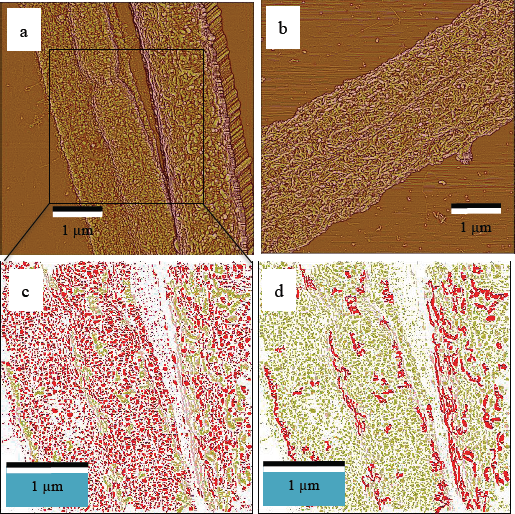
(a) is a high pass filtered AFM images of both immature, collapsed sheath with cell inside (left hand side) and a mature intact sheath (right hand side). In an extreme case (b) exceptionally long fibers parallel to the length of the tube have been observed. Grain size analysis of the boxed region in (a) is detailed in (c) and (d). The analysis on the same region indicates that the younger fresh sheath is primarily made of smaller fibrils (c, red spots) and the older sheath has more long fibrils (d, red squiggly lines) indicating a difference between the two phases of sheath growth. The older sheath (b) is almost exclusively made of long fibrils (analysis not shown but obviously visible in the image.) Figures 4a,b color scale represents height out of the plane of the paper that is 0nm (dark brown) to 50nm (white) for all images. Figures 4c,d identifies particles of different sizes.

**Figure 5:**
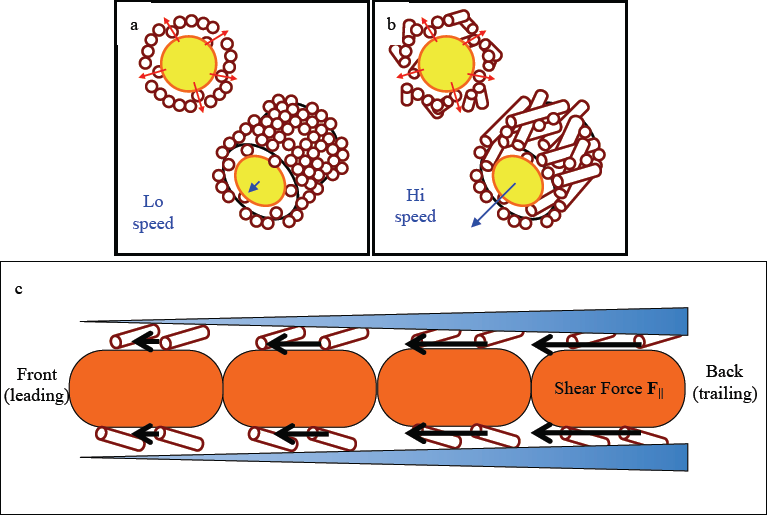
Based on Figure 4's different grain sizes of the sheath materials we speculate the following. In (a) the fibrils within the sheath are globular perhaps do to low speed cell motility, whereas in (b) the fibrils are much longer because the cells are moving at higher speed along the length of the developing sheath. Further Figure 3 suggests a mechanism for growth of the sheath that is described in (c): a proposed motility mechanism based on extrusion of sheath material that pushes the cell train forward through an average shear force ″**F**ǁ″ as the sheath material thickens and hardens behind the advancing cell train. The shear force is greatest where the sheath squeezes tighter, and less at the leading edge (black arrows).

## Discussion

### Sheath Characterization

The highest resolution AFM imaging comes from imaging dried samples, a process that causes some sheaths to collapse, seen in Figure 1. We will use capillary forces to estimate a mechanical property associated with the sheaths. The Young-LaPlace Equation can be used to estimate the capillary pressure on a drying sheath due to surface tension:

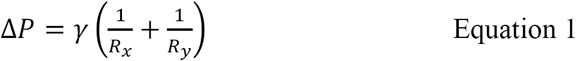

Where γ is the surface tension of water (73.39 dyn/cm = 0.07339 N/m at 15°C), and R_x_ and R_y_ are the radius of curvatures both perpendicular and parallel to the sheath. For a long straight sheath R_x_ = ∞ (parallel to the sheath direction) and R_y_ = 600nm = 6×10^-7^m, then the capillary pressure is about 1.2×10^5^ Pascals (> one atmosphere) or 120nN over a square micron of sheath surface. This estimate of the force needed to collapse sheaths by drying is within the range that the AFM probe can manipulate them. We can estimate this force as follows: The sharp probes used in this study (radius of curvature initially ≤5nm) have a spring constant ″k″ of about 10nN/nm. A Hooke's Law calculation of the normal force on the sample in contact mode with a vertical displacement ″Δy″ of 100nm yields force of F=kΔy ≈ 1000nN. The coefficient of friction between mica and silicon tip is 0.07 (Putnam et al. 1995). Thus the shear force in contact AFM imaging mode is on the order of 100nN. Indeed, we have been able to cut the sheaths with sufficient normal force (images not shown).

An important related mechanical property is the elastic modulus (a.k.a. Young's Modulus "Y") that we will now estimate based on AFM image data. Figure 1 shows evidence that undecorated sheaths collapsed upon drying while older sheaths are more robust, presumably from greater structural integrity due to increased mineralization. From the capillary pressure estimate ((Equation 1) and a reasonable assumption of strain (≈ 1%) on the sheath we can approximate the Elastic Modulus *(Y)* of the fresh, non-mineralized sheaths:

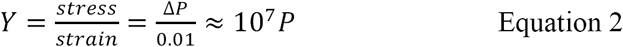

This low value is on the same order of elasticity as rubber, and 100 times smaller than bacteriophage capsids (Ivanovska 2004). The implication is that non-mineralized sheaths are only stable when hydrated and that mineralized sheaths are stiffer, thus do not collapse upon dehydration. AFM techniques have the ability to ″tap″ with large amplitudes and well-characterized forces in order to measure elastic moduli. Future efforts are directed at making elastic modulus measurements to enable us to better characterize the elasticity of the sheaths at different stages in their growth cycle.

The data from Table 1 was generated after making corrections for the finite tip geometry of the AFM probe. Vesenka et al. (1992, 1993) developed simple geometric corrections based on a tube of apparent width ″W_app_″ lying flat on a substrate. The topographic features are a result of artificially broadening (hence ″apparent″) by the finite half angle ″α″ of a pyramidal shaped tip. The resulting corrected diameter ″D″ of the sheath is approximately equal to:

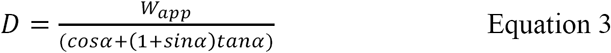

or W=1600nm and *α* =27.5° for the probes used in this study, the apparent diameter of a sheath on the substrate should be about 1000nm.

A collapsed sheath of width ″W″ and height ″H″ above the substrate (double wall thickness of sheath) will have a diameter "D" approximately equal to:

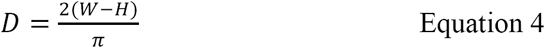

For W = 2200 nm sheath of H = 200 nm, D = 1100 nm, in agreement with the average sheath diameters measured in this study. This confirms the observations that the sheaths are likely composed of a single layer of fibrils described in Figure 2.

### Sheath formation

At present, the best models for sheath formation among the *Leptothrix/Sphaerotilus* family of bacteria come from studies on the heterotrophic *L. cholodnii.* Elegant work by Takeda and colleagues used microslides to grow either *L. cholodnii* (2012a) or *S. natans* (2012b), and directly observe sheath formation. The cell filaments remained stationary, and it was clearly shown that cell growth and elongation and sheath formation co-occurred at the terminus of cell filaments for both species. They proposed that sulfhydryl containing microfibrils are excreted from the cell and diffuse to the sheath layer where they coalesce into a cohesive sheath. It is not known if *L. ochracea* produces similar glycoconjugates. Attempts at staining *L. ochracea* sheaths with fluorescein-labeled N-malemide that binds specifically to free sulfhydryl groups have not been successful (D. Emerson, unpublished results). For these reasons, and other reasons explained below, we believe sheath genesis in *L. ochracea* follows a different model.

There are other important physiological differences between *L. ochracea* and the heterotrophic sheath-formers. *L. ochracea* requires Fe(II) and Fe-oxidation for growth, thus sheath formation and mineral formation appear tightly coupled (Fleming et al., 2011). A recent study of intact, *L. ochracea* mats revealed that empty sheaths formed the bulk of the centimeter-scale mat matrix; *L. ochracea* cells were only present in a 50-100 μm zone at the leading edge of the mat. Time-lapse video (see videos 4,5, & 6 in Chan et al, 2016) of *L. ochracea* cells growing in a capillary tube clearly showed a filament of 30 - 40 cells translocating through liquid medium. There was no visual evidence for an organized sheath at the leading end of the filament, but a fully formed sheath was present at the trailing end of the filament. Presumably the intact sheath was the equivalent of the collapsed, immature or proto-sheaths seen in this work. The length of the cell filament itself did not grow during the short observational period (15 – 30 minutes) indicating sheath formation was dissociated from cell growth. This is fundamentally different from the mechanism of cell growth-based sheath-formation in *L. cholodnii* or *S. natans* as proposed by Takeda et al. (2009) What’s also remarkable about the video observation of Chan et al, (2016) is that the cell filament is moving through the aqueous phase, i.e. not gliding on a surface, with no visible means of propulsion. This suggests an intriguing possibility that *L. ochracea* can propel itself through extrusion of the sheath.

The mostly random orientation of the fibrils in young sheaths, and the estimated small value of elastic modulus ((Equation 2) suggest the fibrils are flexible and sticky (Figures 2 & 3). As evidenced by the width of the fibers (25±5nm) equal to the thickness of the thinnest of immature sheaths (22±3nm), it appears the sheath wall can consist of as little as one monolayer of fibers. Older sheaths with cells inside are thicker, suggesting the fibers become a woven fabric with tensegrity that is maintained by hydration (when the sheaths dry out they collapse). That is older sheaths, most often empty, that continue to oxidize Fe(II) have a more rigid fabric that can resist capillary forces, so that during drying the sheath no longer collapses, and breakage actually causes it to shatter. Presumably the sheath fabric is largely self-organized, otherwise the cell would need to invest significant energy to organize the structure.

#### Model of sheath formation

Our results suggest an initial model for how *L. ochracea* produces an organized sheath._We speculate that orientation of the extruded fibrils might be driven by a combination of random diffusion and oriented filament motion (Figure 5a,b). Diffusion of the cells laterally and rotationally alone by the aqueous medium could explain random fibril orientations with respect to the sheath axis. Single cells are on a size scale (< 1 |im) such that diffusion-driven motion can be substantial. If we estimate their diameter as a hydrodynamic radius, R_h_, of 400nm, this would result in a lateral diffusion constant of:

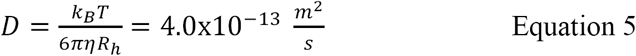

Where k_B_ is Boltzmann's constant, ″T″ is temperature in Kelvin, and ″*η*″ is the viscosity of the medium.

Bacterium of this size can travel a distance of the two dimensional sheath width on the order of a second (Δt = <R^2^>/4D), where Δt is time interval, <R^2^> is the root mean square hydrodynamic radius and D is defined in (Equation 2. This time scale is insufficient for a scaffold to be established. However, almost all of the cells observed in this study appeared to be connected end-to-end during the growth stage of sheaths in a filament. This increases the hydrodynamic radius to micrometer length scales and thus reduces the diffusion coefficient (and decreases the diffusion time) by one or two orders of magnitude. In the absence of any other fluid dynamic interaction the smaller diffusion constant in a chain of cells would allow the scaffold to develop in a globular and random fashion as seen in Figure 4a, and schematically sketched in Figure 5a. Next to the immature sheath ( Figure 4a, at left with cells inside) is a mature sheath with long fibers parallel to the direction of the sheath length at right in the image. It is likely that mineralization is taking place all the time, so the grains grow in size as it appeared in the mature sheath and grain size analysis in Figure 4d. The image shows that there does not appear a preferred direction of orientation of the fibrils. However, Figure 4b is of a mature sheath with a very definite orientation of fibrils parallel to the sheath. We speculate in Figure 5a that slow growth yields more random orientation fibrils and Figure 5b might be explained by more rapid sheath growth. We speculate that the fibril orientation may be a result of lateral motion of the cell filaments during sheath production.

Since the images suggest a preferred orientation of the fibrils parallel or diagonal to the long length of the sheaths (Figures 4b), but not perpendicular, there must be some kind of mechanism that allows lateral motion of the cell filaments in the sheath, though no evidence of flagella have yet been observed using AFM. Recent video analysis (Chan et al., 2016) indicates the cells within sheaths grow parallel to oxygen and dissolved iron gradients. The resolution is insufficient to identify molecular motors pushing them along, but we can rule out hydrodynamic effects since the growth took place in static capillary tubes without the assistance of fluid flow. Based on the evidence collected in Figure 3 we speculate that the mechanism for motion is due to exuding sheath material. As the sheath material becomes increasingly dense due to the combinatorial effects of additional sheath material being added to the sheath, and Fe-oxides forming on the sheath, this results in a shear force that pushes the end of the cell line forward as described below.

#### Model for cell motility

There are several known mechanisms of bacterial motility (Jarrell and McBride, 2008). The most common and well studied is cell swimming aided by flagella, either external, or in the case of spirochetes, intracellular. Gliding along surfaces is another common means of locomotion that is found in diverse bacteria that possess different gliding mechanisms. Another form of translocation is pili-based twitching motility that allows cells to move smaller distances over surfaces. There are also bacteria that lack flagellar, yet are able to swim, the best studied example being the cyanobacterium *Synechococcus* (Ehlers and Oster, 2012).

In what appears to be a novel form of motility, filaments of *L. ohcracea* cells might propel themselves through aqueous media via sheath production. Figure 5c proposes a model for how this motility might occur based on the observed extrusion of sheath material shown in Figures 3a-c. One possibility *we will rule out,* but is valuable for reference purposes, is that the cell motion is driven by a pressure gradient between opposite ends of a sheath. The pressure gradient is caused by the resistance of the cells moving through the sheath, just like the viscosity of blood is responsible for blood pressure between the heart and the rest of the circulatory system. Under these conditions the Hagen-Poiseuille relationship could apply. Calculations will show that this externally driven flow will prove to be insufficient to drive cell motion, but provides an important reference point in terms of a pressure difference between opposite ends of the sheaths. We then show that cell motility can be explained in terms of an *internal* pressure gradient driven by sheath production of the cells themselves.

The experimental speed ″v″ of sheath formation found previously in in laboratory measurements was 27mm/day=0.30μm/s (Chan et al 2016). We use this value also as an estimate for the speed of cell train motion along the newly forming sheath as well, since the vast majority of cells are found on the leading growth edge of the sheath. For a characteristic sheath diameter ″d″ = 2R of about 1μm this corresponds to a Reynolds number of:

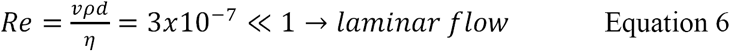

where ″ρ″ = density of water of 1000kg/m^3^ and ″*η*″ is the viscosity of water ≈ 0.0010 Pa*s at ambient temperatures. For laminar flow we can estimate the pressure difference ″ΔP″ from back to front of the cell train using the Hagen-Poiseuille relationship.

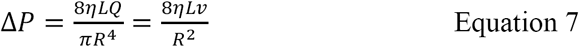

Where Q is the flow rate of the cells equal πR^2^v, the cross sectional area of the sheath times the cells' speed through the sheath, ″L″ is the length scale of a typical chain of cells (≈20 cells at 3μm/cell = 60μm) and the average sheath radius is taken from Table 1 (0.5 μm). This calculation yields a pressure difference across the length of cells of about 0.60 Pa. Over the cross sectional area ″a″=πr^2^ of a cell of radius ″r″ the normal force F=ΔP*a = 3.6×10^-14^N, which is an insignificant force on this microscopic scale and is probably not responsible for pushing the line of cells (cell ″train″) forward.

However, if we speculate the force comes from *shear* interactions from *within* the sheath as the scaffold material is extruded along the length of the cell chain and squeezing the cells forward against the thickening and hardening sheath (Figure 5c), we can assume an area of the sheath's cylindrical shape along the length of the cell chain A = 2πr*L, resulting in a shear force F=ΔP*A = 4.5×10^-10^N, or 0.45nN, which is about 40x larger than the force exerted by a bacterial flagellum (e.g. Darnton and Berg, 2006). This would cause the cell filament to translocate in a completely aqueous medium and could account for the motility observed by Chan et al. (2016).

We are not aware of any other reports of this type of microbial shear force based propulsion that allow advancement and movement of a bacterial cell filament in the purely aqueous phase. An additional fascinating aspect of this motility is that it is also coupled to directionality in intact microbial mats formed by *L. ochracea.* The recent paper by Chan et al (2016) showed that *L. ochracea* sheaths are able to move in gradients of Fe(II) and O_2_ in a uniform direction, i.e. nonrandomly, indicating there is a link between the cells ability to couple chemo-sensing with motility. The mechanism for this process remains unknown.

## Acknowledgements

Acknowledgements.We would like to thank Dr. Emily Fleming for her help in sample collection and preparation, as well as helpful discussions. DE was supported by the National Science Foundation IOS 0951077 and NASA Exobiology grant NNX15AM11G. We also acknowledge receipt of an NSF Research Opportunity Award to support work in JV’s laboratory at UNE.

